# Fenbendazole controls *in vitro* growth, virulence potential and animal infection in the *Cryptococcus* model

**DOI:** 10.1101/2020.02.13.948745

**Authors:** Haroldo C. de Oliveira, Luna S. Joffe, Karina S. Simon, Rafael F. Castelli, Flavia C. G. Reis, Arielle M. Bryan, Beatriz S. Borges, Lia C. Soares Medeiros, Anamelia L. Bocca, Maurizio Del Poeta, Marcio L. Rodrigues

## Abstract

The human diseases caused by the fungal pathogens *Cryptococcus neoformans* and *C. gattii* are associated with high indices of mortality, and toxic and/or cost-prohibitive therapeutic protocols. The need for affordable antifungals to combat cryptococcal disease is unquestionable. Previous studies suggested benzimidazoles as promising anti-cryptococcal agents combining low cost and high antifungal efficacy, but their therapeutic potential has not been demonstrated so far. In this study, we investigated the antifungal potential of fenbendazole, the most effective anti-cryptococcal benzimidazole. Fenbendazole was inhibitory against 30 different isolates of *C. neoformans* and *C. gattii* at a low concentration. The mechanism of anti-cryptococcal activity of fenbendazole involved microtubule disorganization, as previously described for human parasites. In combination with fenbendazole, the concentrations of the standard antifungal amphotericin B required to control cryptococcal growth were lower than those required when this antifungal was used alone. Fenbendazole was not toxic to mammalian cells. During macrophage infection, the anti-cryptococcal effects of fenbendazole included inhibition of intracellular proliferation rates and reduced phagocytic escape through vomocytosis. Fenbendazole deeply affected the cryptococcal capsule. In a mice model of cryptococcosis, the efficacy of fenbendazole to control animal mortality was similar to that observed for amphotericin B. These results indicate that fenbendazole is a promising candidate for the future development of an efficient and affordable therapeutic tool to combat cryptococcosis.

## Introduction

Cryptococcosis caused by *Cryptococcus neoformans* and *C. gattii* kills almost 200,000 humans each year (1). The disease, which is devastating in sub-Saharan Africa, significantly affects other regions of the globe, including Asia, Oceania, Europe and the Americas (2–4). The World Health Organization (WHO) recommends three therapeutic phases for treating cryptococcal meningitis, including an induction therapy with amphotericin B plus flucytosine (week 1) followed by fluconazole (week 2), a consolidation phase with fluconazole (weeks 3-10) and maintenance therapy of up to 12 months also with fluconazole (5). However, amphotericin B and flucytosine are not available in many countries (6). In addition, the effective treatment of human cryptococcosis is cost-prohibitive in most of the regions that are severely affected by *C. neoformans* and *C. gattii*. In Malawi, Zambia, Cameroon, and Tanzania, the costs for treating each patient with cryptococcal meningitis includes US $1442 for 2 weeks of oral fluconazole and flucytosine, $1763 for 1 week of amphotericin B and fluconazole, $1861 for 1 week of amphotericin B and flucytosine, $2125 for 2 weeks of amphotericin B and fluconazole, and $2285 for 2 weeks of amphotericin B and flucytosine (7). In Brazil, the therapeutic costs of lipid formulations of amphotericin B can exceed US $100,000 per patient (8). In summary, novel therapeutic protocols for treating the diseases caused by *C. neoformans* and *C. gattii* are urgent. However, the development of novel drugs is time-consuming, highly expensive and commonly unsuccessful (9, 10). In this context, drug repurposing has emerged as a promising alternative for the development of novel therapies against neglected pathogens, including fungi (11–13).

*C. neoformans* and *C. gattii* are sensitive to benzimidazoles *in vitro* (14). Flubendazole is inhibitory against all pathogenic *Cryptococcus* species, including isolates that are resistant to fluconazole (13). In mice, orally administered flubendazole resulted in an expressive reduction in fungal burden in comparison with controls, but in the rabbit model, there were no quantifiable drug concentrations or antifungal activity in the cerebrospinal fluid or brain (15). Mebendazole, another member of the family of benzimidazoles, also showed antifungal activity against *C. neoformans*, including phagocytized yeast cells and cryptococcal biofilms (16).

The anti-cryptococcal effects of other benzimidazoles have been superficially examined. In comparison to other benzimidazoles, fenbendazole was the most efficient compound showing *in vitro* fungicidal activity (16), but mechanistic approaches and *in vivo* activity of this compound were not evaluated. Fenbendazole has been licensed worldwide for the treatment and control of helminth infections in food and non-food producing animals for more than 30 years and its safety is well established. According to the European Medicines Agency (17), fenbendazole had negligible acute toxicity in single-dose animal studies and no points of concern relevant for the safety of fenbendazole in humans could be identified. No treatment-related effects were observed in the offspring of dogs, pigs, sheep, and cattle administered fenbendazole at various times during gestation. Finally, the compound was not genotoxic, and no evidence for carcinogenicity was found (17, 18).

Based on the anti-cryptococcal effects of fenbendazole (16) and negligible toxicity to humans and animals (17, 18), we evaluated the therapeutic potential of this benzimidazole against pathogenic cryptococci. Our results demonstrated that fenbendazole was inhibitory against several strains of *C. neoformans* and *C. gattii*. The mechanism of antifungal activity of fenbendazole involved the functionality of microtubules. Fenbendazole had low toxicity to mammalian cells alone or in combination with amphotericin B and its antifungal effects included inhibition of virulence determinants and reduced proliferation of *Cryptococcus* inside macrophages. Finally, fenbendazole was highly effective in a mice model of cryptococcosis. These results support the use of fenbendazole as a prototype for the development of novel pharmaceutical preparations for treating cryptococcosis.

## Results

### Fenbendazole affects several isolates of *Cryptococcus* and has low toxicity to mammalian cells

We first determined the minimum inhibitory concentration (MIC) of fenbendazole against strains H99 and R265, the standard isolates of *C. neoformans* and *C. gattii*, respectively. A similar MIC of 0.012 μg/ml was found for both strains (Figure 1A). We tested 15 additional isolates of *C. neoformans* and 12 other isolates of *C. gattii* and the same MIC of 0.012 μg/ml was found for all of them, despite their differential susceptibility to fluconazole and amphotericin B (Table 1). At the MIC, fenbendazole exerted fungicidal effects, as concluded from the highly reduced detection of colony forming units (CFU) of *C. neoformans* and *C. gattii* after exposure to the drug (Figure 1B). In a toxicity model using mammalian macrophages, the concentration of fenbendazole required to kill 50% of the cell population (LD50) corresponded to 3.072, generating a selectivity index (LD50/MIC) of 256. This result confirmed the low toxicity of fenbendazole.

**Figure 1.**
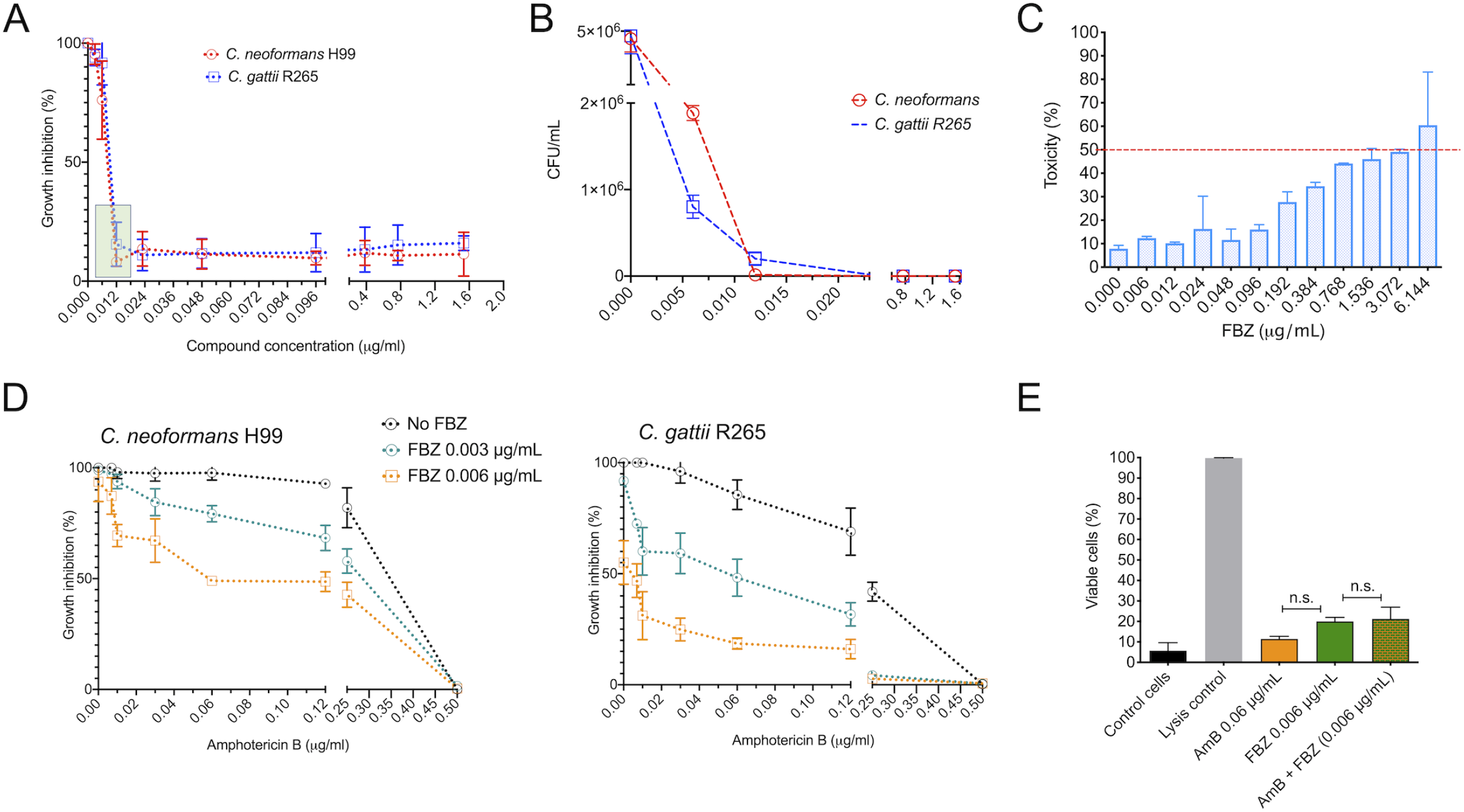
Antifungal effect and toxicity of fenbendazole (FBZ). A. Determination of the minimum inhibitory concentration (MIC) of fenbendazole against *C. neoformans* (strain H99) and *C. gattii* (strain R265). Both strains were sensitive to a MIC of 0.012 μg/ml (boxed area). B. FBZ is fungicidal, as concluded from its ability to drastically reduce the numbers of colony forming units (CFU) of *C. neoformans* and *C. gattii* at the MIC. C. Dose-dependent profile of toxicity of fenbendazole against RAW 264.7 macrophages. The dotted line represents the 50% cut off of cellular viability. D. Antifungal effects of subinhibitory doses of fenbendazole (0.003 and 0.006 μg/ml) in combination with variable concentrations of amphotericin B (AmB) against *C. neoformans* and *C. gattii*. In both cases, the presence of fenbendazole results in decreased concentrations of amphotericin B required for growth inhibition. E. Analysis of the potential of fenbendazole and amphotericin B alone or in combination to kill RAW 264.7 macrophages. Control cells consisted of mammalian cultures treated with vehicle (DMSO) alone. Lysis control consisted of cells treated with 1x lysis solution (provided by the manufacturer). No significant (n.s.) differences were found between the antifungal preparations.

**Table 1.** Determination of MICs (μg/ml) in different isolates of *C. neoformans* and *C. gattii* using fenbendazole (FBZ), fluconazole (FCZ) and amphotericin B as antifungals^1^

| Isolate<sup>2</sup> | MIC |  |  |
| --- | --- | --- | --- |
|  | FBZ | FCZ | AmB |
| *C. neoformans* H99 | 0.012 | 2.0 | 0.5 |
| *C. neoformans* 162 | 0.012 | 8.0 | 0.5 |
| *C. neoformans* 191 | 0.012 | 8.0 | 0.25 |
| *C. neoformans* 186 | 0.012 | 4.0 | 0.25 |
| *C. neoformans* 161 | 0.012 | 4.0 | 0.5 |
| *C. neoformans* 160 | 0.012 | 4.0 | 0.5 |
| *C. neoformans* 139 | 0.012 | 4.0 | 0.5 |
| *C. neoformans* 118 | 0.012 | 8.0 | 0.25 |
| *C. neoformans* 116 | 0.012 | 8.0 | 0.5 |
| *C. neoformans* 115 | 0.012 | 4.0 | 0.5 |
| *C. neoformans* 222 | 0.012 | 2.0 | 0.5 |
| *C. neoformans* 223 | 0.012 | 4.0 | 0.5 |
| *C. neoformans* 225 | 0.012 | 8.0 | 0.5 |
| *C. neoformans* 271 | 0.012 | 4.0 | 0.5 |
| *C. neoformans* 216 | 0.012 | 4.0 | 0.25 |
| *C. neoformans* 218 | 0.012 | 2.0 | 0.25 |
| *C. gattii* R265 | 0.012 | 8.0 | 0.5 |
| *C. gattii* 456 | 0.012 | 2.0 | 0.12 |
| *C. gattii* 460 | 0.012 | 16.0 | 0.25 |
| *C. gattii* 461 | 0.012 | 8.0 | 0.25 |
| *C. gattii* 367 | 0.012 | 8.0 | 0.25 |
| *C. gattii* 368 | 0.012 | 16.0 | 0.25 |
| *C. gattii* 434 | 0.012 | 16.0 | 0.25 |
| *C. gattii* 221 | 0.012 | 8.0 | 0.25 |
| *C. gattii* 365 | 0.012 | 4.0 | 0.5 |
| *C. gattii* 366 | 0.012 | 8.0 | 0.5 |
| *C. gattii* 158 | 0.012 | 4.0 | 0.25 |
| *C. gattii* 188 | 0.012 | 4.0 | 0.5 |
| *C. gattii* 198 | 0.012 | 2.0 | 0.25 |
| *C. krusei* ATCC6258<sup>2</sup> | >1.536 | >64 | 0.25 |
| *C. parapsilosis* ATCC22019<sup>3</sup> | >1.536 | 1.0 | 0.25 |

<sup>1</sup>As preconized by the EUCAST, growth inhibition measurements corresponded to 50% for FLZ and 90% for AmB. FBZ inhibition rates corresponded to 90%.
<sup>2</sup>Strain numbers were attributed by the Collection of Pathogenic Fungi of Fiocruz. For details, see http://cfp.fiocruz.br/index.
<sup>3,4</sup>The *C. krusei* and *C. parapsilosis* isolates were used as controls of antifungal activity, as preconized by the EUCAST. The MIC results obtained with these isolates were at the expected range (39).

Fenbendazole potentiated the antifungal activity of amphotericin B, as inferred from the results obtained from the checkerboard assay. When used alone, amphotericin B produced a MIC of 0.5 μg/ml against both *C. neoformans* (H99 strain) and *C. gattii* (R265 strain). However, when tested in combination with 0.006 μg/ml fenbendazole, the amphotericin B MIC was 8-fold decreased (0.06 μg/ml). Indeed, dose-response analyses efficiently illustrate that the anti-cryptococcal effects of amphotericin B are boosted by different concentrations of fenbendazole, especially against *C. gattii* (Figure 1D). Analysis of the toxic effects of the compounds in combination or alone revealed that all systems had low and similar toxicity to mammalian macrophages (Figure 1E). The fractional inhibitory concentration index (FICI) corresponded to 0.62 for the H99 strain and 0.49 for the R265 isolate, suggesting that fenbendazole was slightly more synergistic with amphotericin B against *C. gattii* than *C. neoformans*.

### Fenbendazole affects β-tubulin distribution in *C. neoformans* and *C. gattii*

We asked whether the mechanism of anti-cryptococcal activity of fenbendazole was similar to what was previously described for *C. gattii* (16) or if it was related to its well characterized anthelmintic effect (19–21). Mutant strains of *C. gattii* lacking expression of the Aim25 scramblase or the nucleolar protein Nop16 were resistant to mebendazole (16), suggesting that these proteins are potential targets for the antifungal activity of benzimidazoles. To investigate the mechanism of action of fenbendazole, we first evaluated its antifungal activity against mutant strains of *C. gattii* lacking the *AIM25* or *NOP16* genes and observed that both strains and wild type cells were similarly sensitive to this benzimidazole (Figures 2A and B). This result suggested that, in contrast to mebendazole, Aim25 and Nop16 are not required for the antifungal activity of fenbendazole. We then asked whether the inhibitory effect of fenbendazole against *C. neoformans* and *C. gattii* involves interference with the functions of β-tubulin, as consistently described for parasites (19, 21). Staining of β-tubulin in fungal cells revealed that fenbendazole-treated fungi and untreated cryptococci had markedly different profiles of microtubule organization. Control cells showed high fluorescence intensity and a well-defined intracellular pattern of β-tubulin staining (Figure 2C). Fluorescence detection was apparently less intense in drug-treated cells, which also manifested a markedly more disperse staining of β-tubulin. In addition, the effects of fenbendazole on β-tubulin staining were apparently more drastic in *C. gattii* than in *C. neoformans*, as concluded from the weaker signals of β-tubulin staining in the former species. These results indicate that the mechanism antifungal activity of fenbendazole against *Cryptococcus* is similar to that described for human parasites.

**Figure 2.**
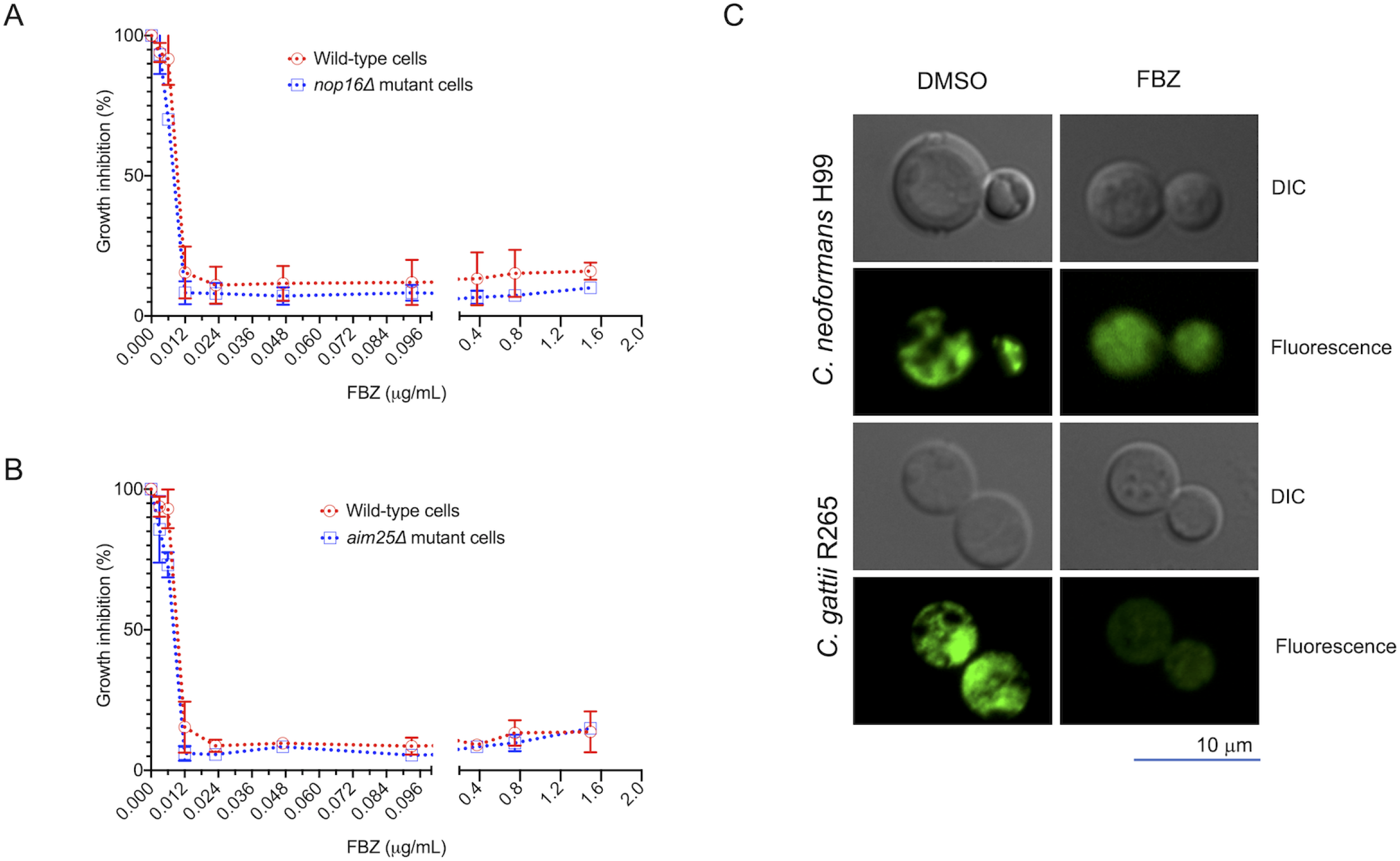
The antifungal effects of fenbendazole are related to microtubule disorganization. Nop16 and Aim25, which were previously suggested to be the targets of the antifungal activity of benzimidazoles in *C. gattii*, are not involved in the antifungal activity of fenbendazole (FBZ), as concluded from the similar growth rates observed for wild-type *C. gattii* and mutant cells lacking *NOP16* (A) or *AIM25* (B) in the presence of variable concentrations of FBZ. A comparison between the β-tubulin staining pattern in control cells (DMSO) and FBZ-treated fungi (C) revealed that drug treatment profoundly affected microtubule distribution, with apparently more intensive effects on *C. gattii*. Fungal cells are shown under the under differential interference contrast (DIC) and fluorescence modes.

### Intranasal administration of fenbendazole results in the control of animal cryptococcosis

The combination of low toxicity, efficient antifungal efficacy, and defined mechanism of action led us to test the antifungal potential of fenbendazole *in vivo*. To reduce the number of animals used to a minimum, we selected the standard strain H99 of *C. neoformans* for the *in vivo* work, based on the highest prevalence of this species in human disease (3). Metabolization by host tissues is a common feature of orally administered benzimidazoles (22). Fenbendazole, specifically, is rapidly sulfoxidated by liver microsomes after oral absorption (22). Therefore, we first tested the in vitro antifungal activity of the sulfone-derivative of fenbendazole, which showed no inhibitory potential (Figure 3A). We performed similar tests with the liver metabolites of other benzimidazoles (mebendazole, flubendazole and albendazole) and none of them manifested antifungal activity (Figure 3B-D), suggesting that host metabolization is an important limitation for the antifungal activity of benzimidazoles in general. In fact, orally administered fenbendazole and mebendazole had no effects on mice cryptococcosis (data not shown). Therefore, to avoid liver metabolization and to promote an increased bioavailability of fenbendazole in its native form, we administered the drug intranasally. Under these conditions, mice receiving amphotericin B and fenbendazole had similarly high survival rates, in comparison with vehicle-treated mice (P = 0.0014, Figure 3E). The experiment was repeated under the same conditions and identical results were obtained.

**Figure 3.**
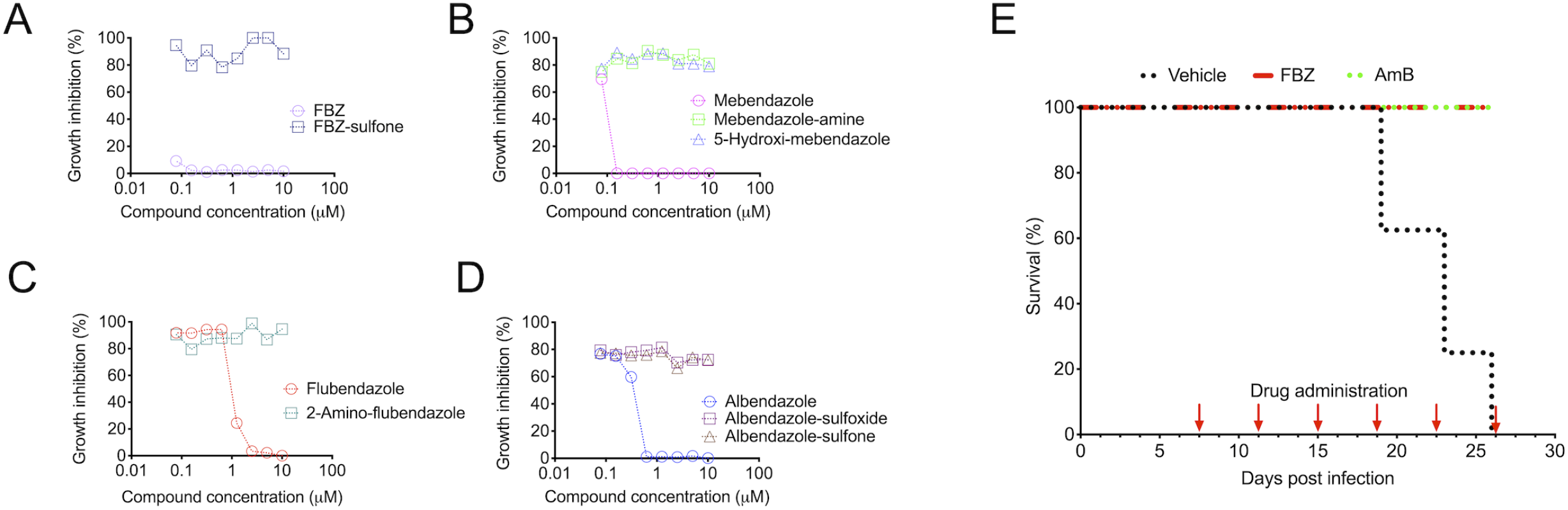
Analysis of the potential of fenbendazole to control animal cryptococcosis. A-D. Antifungal effects of fenbendazole and other benzimidazoles in comparison with their liver metabolites. All benzimidazoles had clear antifungal activity, in contrast to their liver metabolites of fenbendazole (A), mebendazole (B), flubendazole (C) and albendazole (D). E. Treatment of lethally infected mice with intranasally delivered fenbendazole or intraperitoneally administered amphotericin B (AmB). All vehicle-treated animals died 26 days post-infection. Drug-treated animals were all alive 27 days post infection.

### Fenbendazole affects the virulence potential of *Cryptococcus*

We asked whether the high efficacy of fenbendazole *in vivo* was related to neutralization of virulence determinants in addition to its antifungal effects. Capsule synthesis and intracellular proliferation rates have been consistently associated to the pathogenic potential of cryptococci (23, 24). We therefore evaluated whether fenbendazole was able to interfere with each of these biological events.

Microscopic analysis of fenbendazole-treated *C. neoformans* and *C. gattii* revealed clear effects on the capsular architecture, although some species-specific particularities were observed. In all cases, fungal aggregates with reduced capsular dimensions were observed after drug treatment (Figure 4A). Scanning electron microscopy of the H99 strain of *C. neoformans* after exposure to fenbendazole revealed shorter and scarcer capsular fibers, in comparison with control cells. This perception was confirmed by immunofluorescence analysis. In *C. gattii* (strain R265), surface fibers and capsular dimensions were also reduced after exposure to fenbendazole. However, in comparison with drug-treated *C. neoformans*, fenbendazole was apparently less efficient in reducing the capsular dimensions of *C. gattii*. In both cases, a dose-dependent reduction of capsular dimensions was observed after cryptococci were treated with fenbendazole (Figure 4B).

**Figure 4.**
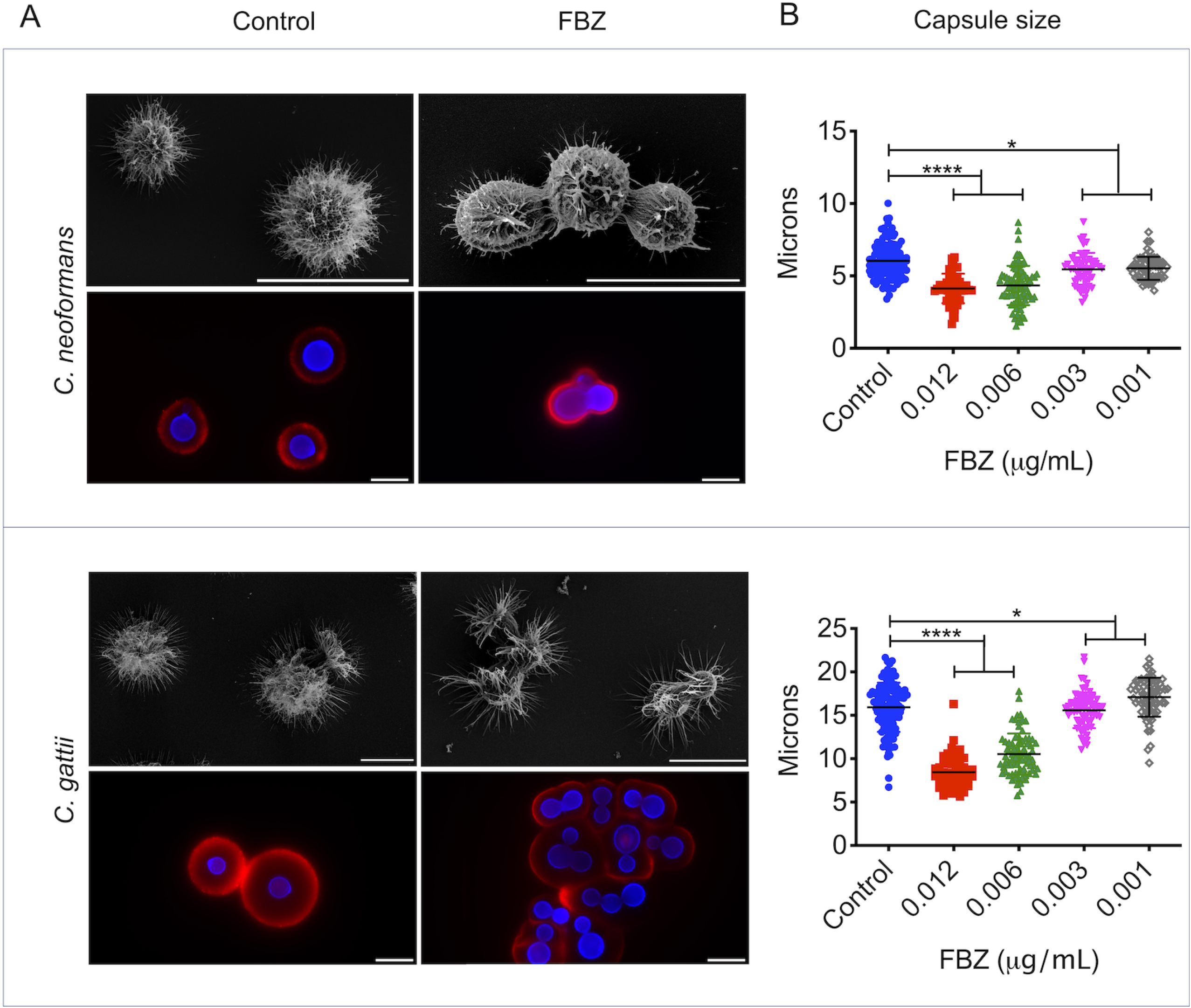
Fenbendazole (FBZ) affects capsule formation in *C. neoformans* and *C. gattii*. A. Morphological analysis of vehicle (control) and drug-tread *C. neoformans* and *C. gattii*. For each species, upper panels illustrate capsular alterations by scanning electron microscopy, and lower panels illustrate capsular morphology (red fluorescence) and cell wall staining (blue fluorescence). Scale bars, 10 μm. B. Quantitative determination of capsular dimensions after treatment of fungal cells with vehicle alone (control) or variable concentrations of fenbendazole. Asterisks denote P < 0.05 (*) or P < 0.0001 (****).

Due to the suggested link between therapeutic failure and intracellular proliferation of cryptococci (23, 25, 26), we evaluated whether fenbendazole could influence the fate of *C. neoformans* and *C. gattii* in infected macrophages. Initial microscopic observation suggested that fenbendazole affected the cell division frequency of both *C. neoformans* and *C. gattii* (Figures 5A and B). In macrophages infected with *C. neoformans* that were further treated with DMSO, intracellular division was initially observed 1 h after internalization of fungal cells, with a clear peak at 5 h post phagocytosis (Figure 5C). In drug-treated systems, intracellular proliferation was first observed 4 h after phagocytosis, with less intense peaks at 5 and 7 h post infection. In similar systems using *C. gattii*, intracellular proliferation in vehicle-treated macrophages was first observed 1 h post-infection, with more prominent peaks of replication at 5 and 7 h post-infection (Figure 5D). In drug-treated systems, fungal intracellular proliferation was only observed after 5 h. These results were suggestive of lower intracellular proliferation rates (IPR) of phagocytized *C. neoformans* and *C. gattii*. Since low IPR have been linked with reduced virulence (23, 25, 27), we investigated the effects of fenbendazole on this important parameter.

**Figure 5.**
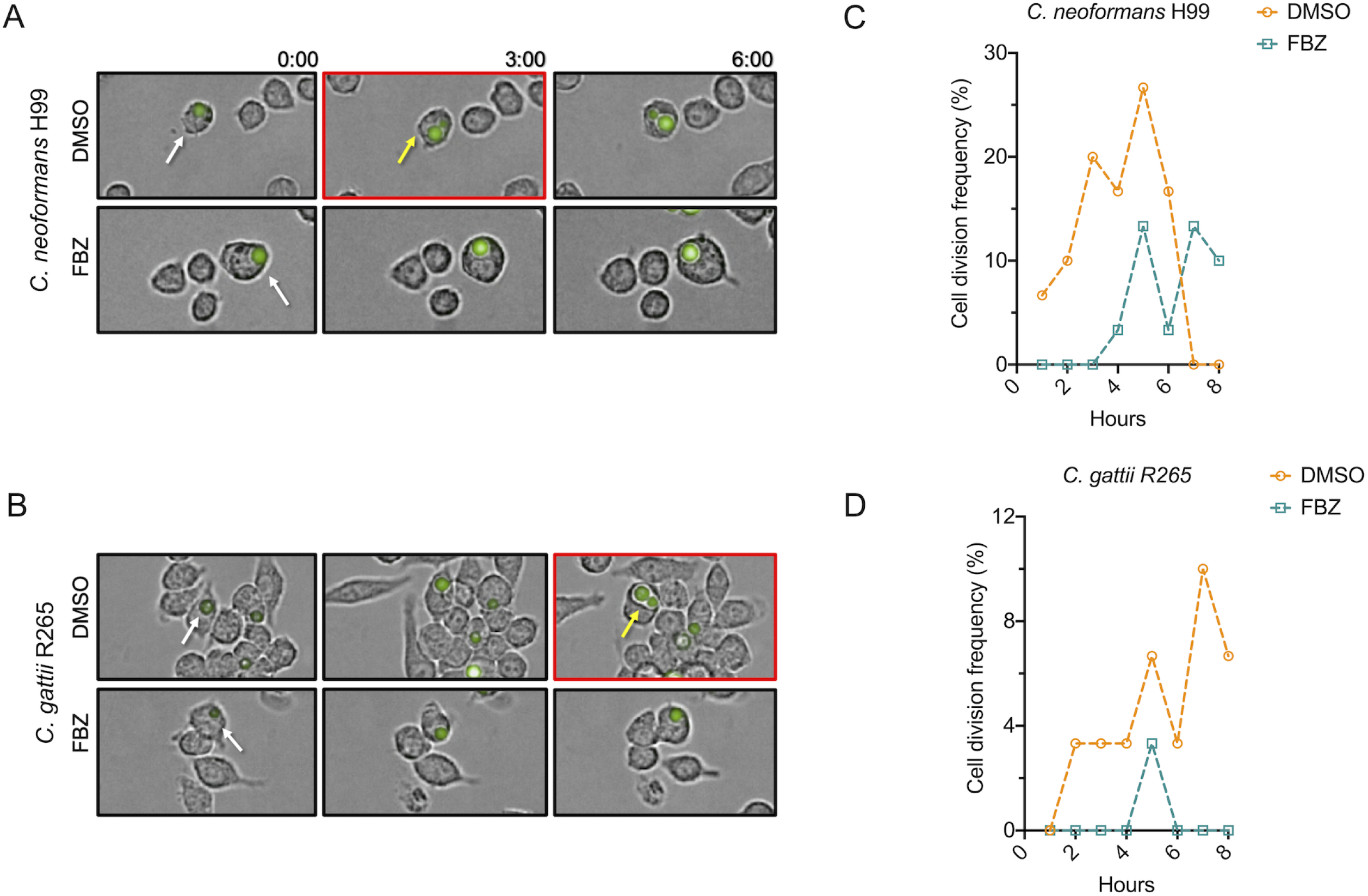
Time-lapse detection of budding yeast cells in macrophages infected with *C. neoformans* (strain H99) or *C. gattii* (strain R265). Video-microscopy analysis of macrophages infected with single-cell *C. neoformans* (A) or *C. gattii* (B) (white arrows) suggested that detection of yeast budding (yellow arrows) is delayed in fenbendazole (FBZ)-treated systems, in comparison with vehicle (DMSO)-treated macrophages. Panels boxed in red denote the initial detection of yeast budding. This initial perception was confirmed by a quantitative analysis of intracellular budding for *C. neoformans* (C) and *C. gattii* (D).

In both species, exposure of infected macrophages to fenbendazole resulted in reduced IPR at both inhibitory and sub inhibitory concentrations (Figures 6A and 6B). High IPR are related to vomocytosis, a mechanism of phagocytic scape commonly used by *Cryptococcus* (28, 29). We therefore determined the vomocytosis levels after exposure of infected macrophages to fenbendazole. Similarly, fenbendazole reduced the rates of vomocytosis at both inhibitory and sub inhibitory concentrations in macrophages infected with *C. neoformans* or *C. gattii* (Figures 6C and D). Since IPR and vomocytosis are directly linked to the ability of *Cryptococcus* to survive after ingestion by phagocytic cells, we also counted viable CFU of *C. neoformans* and *C. gattii* after interaction with the macrophages. Once again, treatment of the mammalian cells with fenbendazole affected the growth of *C. neoformans* and *C. gattii*. In both cases, CFU counts were reduced after treatment of infected cells with fenbendazole (Figures 6E and F). Once again, fenbendazole promoted an increase in the antifungal efficacy of amphotericin B, as inferred from the reduced concentration of this antifungal required for the intracellular killing of *C. neoformans* and *C. gattii* (Figures 6E and F).

**Figure 6.**
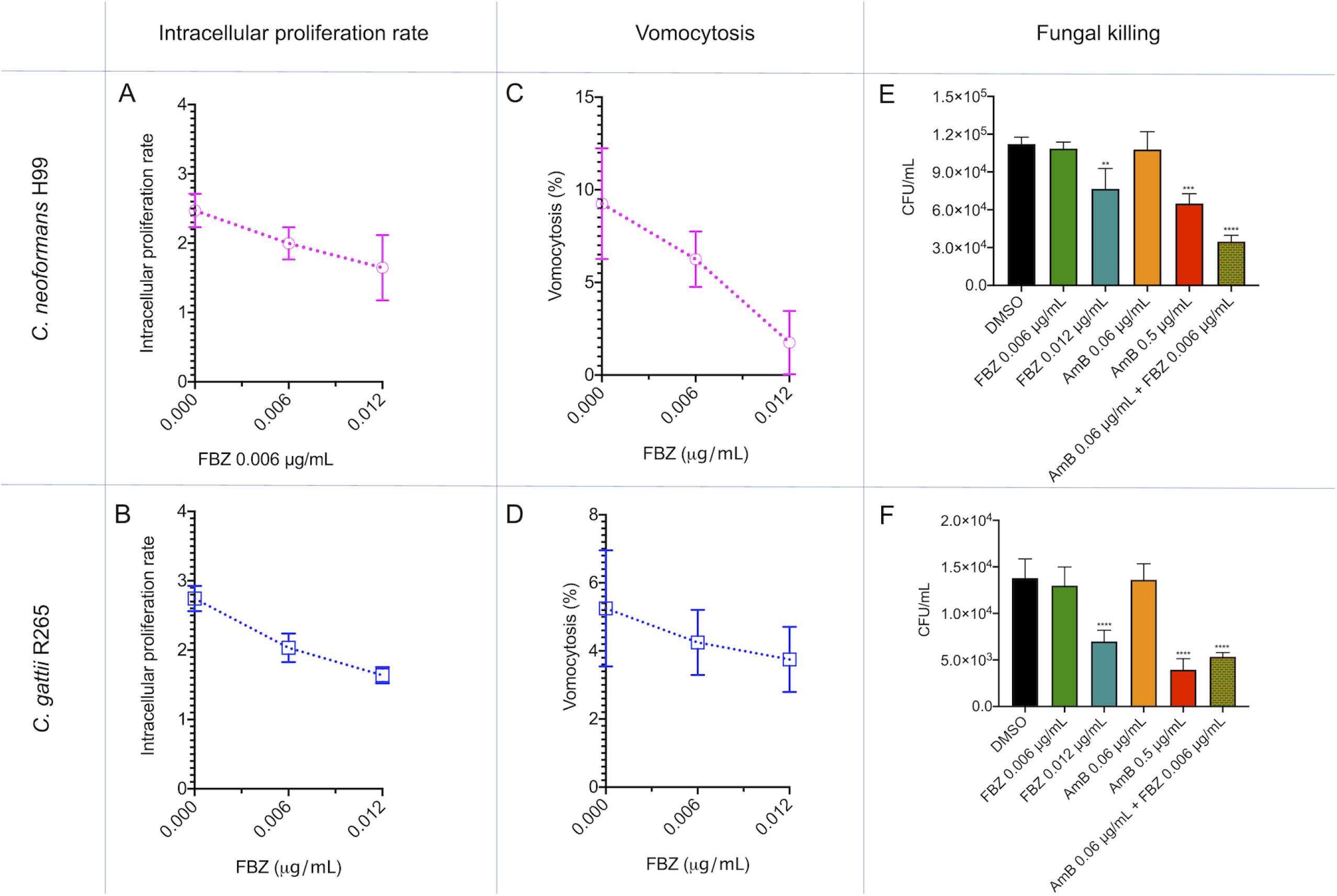
Effects of fenbendazole (FBZ) on the intracellular fate of *Cryptococcus*. Treatment with fenbendazole decreased the intracellular proliferation rates of *C. neoformans* (A) and *C. gattii* (B). Similar profiles of inhibitory effects were observed when vomocytosis was analyzed in macrophages infected with *C. neoformans* (C) or *C. gattii* (D). The recovery of significantly lower viable fungal cells from FBZ-treated macrophages (0.012 μg/ml) indicated intracellular activity against *C. neoformans* (E) and *C. gattii* (F). Similar results were obtained with an antifungal concentration range of amphotericin B (AmB) or with a combination of subinhibitory doses of AmB and FBZ. **P = 0.01; ***P = 0.001; ****P<0.0001.

## Discussion

The *in vitro* anti-cryptococcal activity of benzimidazoles has been consistently reported (13–16). A general analysis of these reports on the anti-cryptococcal activity of benzimidazoles indicated that, among all members of this drug family, fenbendazole required the lowest MIC against *C. neoformans* and *C. gattii*, in comparison to other benzimidazoles (13–16). In addition, a detailed study on the toxicity of fenbendazole suggested high levels of safety in humans (17, 18). In our study, we confirmed the efficacy of fenbendazole as an anti-cryptococcal agent. We tested approximately 30 strains of *C. neoformans* and *C. gattii* and observed that they were all similarly susceptible to fenbendazole, suggesting that resistance to this benzimidazole might not be an issue in pathogenic cryptococci. A similar MIC of 0.012 μg/ml was found for all of them. For comparison, a recent study (15) analyzed 50 strains of *C. neoformans* according to their susceptibility to flubendazole, another member of the benzimidazole family. Using the same EUCAST methodology, MICs of 0.03 (one strain), 0.06 (19 strains), 0.125 (25 strains) and 0.25 (5 strains) μg/ml were found in this study. These values were 2.5 to 20-fold higher than the MIC found for fenbendazole in this study. The low toxicity of fenbendazole was also certified in our essay, as well as its ability to improve the antifungal activity of amphotericin B. These characteristics led us to explore the properties of fenbendazole in mechanistic and therapeutic models.

In a previous study, we used a mutant collection of *C. gattii* for determination of the cellular targets of mebendazole, another anti-cryptococcal benzimidazole (16). These mutants were screened for resistance phenotypes in the presence of mebendazole, based on the assumption that, in the absence of a cellular target required for antifungal activity, the drug would lack anti-cryptococcal properties. The mutants showing highest levels of resistance to mebendazole lacked expression of Aim25, a cryptococcal scramblase (30), or Nop16, a putative nucleolar protein (16). We initially assumed that these targets were required for the activity of fenbendazole, but our current results indicate that they are not involved in the antifungal properties of this benzimidazole. In nematodes, the benzimidazoles bind to β-tubulin, leading to local unfolding of the protein and consequent abnormal conformation. This outcome finally results in the inhibition of the polymerization of α- and β-tubulin subunits to form microtubules, causing lethal effects in dividing cells (31). In the *C. neoformans* model, a similar mechanism was attributed to flubendazole and other benzimidazoles (32). However, fenbendazole was not specifically tested in these studies. In addition, the cellular effects of benzimidazoles on the cryptococcal microtubules were not reported. In our study, treatment of both *C. neoformans* and *C. gattii* with fenbendazole led to a clear alteration in β-tubulin detection, strongly suggesting that its mechanism of action is related to that described for the nematodes and other parasites.

The benzimidazoles are extensively metabolized in mammals following oral administration. The parent compounds are generally short-lived, and metabolites predominate in plasma, tissues and excreta (22). Fenbendazole, for instance, is rapidly metabolized by liver microsomes after oral absorption. In pigs, this benzimidazole was rapidly absorbed after oral administration, but its systemic bioavailability was low (33). The intranasal route, on the other hand, transport drugs directly to the brain from the nasal cavity along the olfactory and trigeminal nerves (34), avoiding the first-pass metabolism in liver and gastrointestinal tract. These findings agree with our current results showing that all animals infected with *C. neoformans* and treated with fenbendazole intranasally survived. However, our results also suggest that liver metabolization is a limitation for the oral treatment of fungal diseases with fenbendazole. In this sense, strategies for protecting drug candidates against metabolic modifications are widely available (35) and they could be applied to the use of fenbendazole as a scaffold for antifungal development. Most likely, intranasal pharmaceutical preparations of fenbendazole could be promising and safe therapeutic alternatives in the *Cryptococcus* model.

Interference with pathogenic traits in addition to primary antifungal effects were likely related to the high efficacy of fenbendazole in the control of animal cryptococcosis observed in our study. High levels of uptake cryptococci by macrophages in vitro were associated with long-term survival of human patients (36). Importantly, strains showing high uptake by macrophages were hypocapsular. The connections between phagocytic events, reduced capsules and the outcome of human disease consisted of the basis for our studies on the effects of fenbendazole on capsular architecture and outcome of macrophages infection. Fenbendazole produced effects that are potentially positive to the host in all cases, since it caused a decrease in capsular density and dimensions, in addition to reducing intracellular proliferation rates and events required for phagocytic escape.

It was estimated that the cost to procure one million doses of standard benzimidazoles (500 mg each) would be approximately US$ 20,000, including international transport (37). In comparison to the cost of treating cryptococcosis patients with lipid formulations of amphotericin B (8, 9), these numbers are extraordinarily low. In this context, our results support the development of pharmaceutical preparations of fenbendazole to be tested as alternative anti-cryptococcal agents. If effective in humans, fenbendazole could represent an affordable alternative for the treatment of a disease that has an extremely negative impact on the health conditions of neglected populations.

## Material and methods

### Strains and Growth Conditions

*Cryptococcus neoformans* and *C. gattii* (strains H99 and R265, respectively) were used in most experiments. In addition, 15 isolates of *C. neoformans* and 12 isolates of *C. gattii*, obtained from the Collection of Pathogenic Fungi available at Fiocruz, were tested for sensibility to fenbendazole. For studies of mechanism of antifungal activity, the *C. gattii* mutant strains *aim25Δ* and *nop16Δ* were also tested (16, 30). For the *in vitro* interaction assays, green fluorescence protein (GFP)-tagged *C. neoformans* and *C. gattii* were used (38). These strains, which were kindly provided by Dr. Robin May, were constructed in the H99 (*C. neoformans*) and R265 (*C. gattii*) backgrounds, respectively. All strains were maintained in Sabouraud agar plates and cultivated in the Sabouraud medium at 30°C for 24 h before all experiments. RAW 264.7 macrophages were used for toxicity and phagocytosis experiments. The cells were maintained in the Dulbecco’s Modified Eagle Medium (DMEM) medium (Sigma Aldrich, catalog number D5796) supplemented with 10% heat inactivated fetal bovine serum (FBS) at 37°C under a 5% CO_2_ atmosphere.

### Antifungal Susceptibility Testing

Fenbendazole was obtained from Sigma Aldrich (catalog number F5396) in its solid form. Stock solutions were prepared in DMSO (drug vehicle). In the antifungal susceptibility tests, vehicle concentration was kept at 1%. To determine the minimal inhibition concentration (MIC) of fenbendazole, the broth microdilution method was used, following the protocols established by the European Committee on Antimicrobial Susceptibility Testing (EUCAST; E.Def 7.3.1 reference method) (39). Plates to be used in the antifungal susceptibility tests contained the Roswell Park Memorial Institute (RPMI) medium supplemented with 2% glucose, buffered with 165 mM 3-(*N*-morpholino)propanesulfonic acid (MOPS; pH 7.0). The medium was supplemented with fenbendazole in the concentration range of 0.003 to 1.536 μg/ml. Control systems contained the drug vehicle alone. Before inoculation in the antifungal testing plates, *C. neoformans* and *C. gattii* were grown on Sabouraud agar plates at 30°C for 48 h. A suspension of 2.5 x 10^5^ cells/ml was prepared in distilled water and 100 μl of this suspension was transferred to each well of 96-plates containing 100 μl of RPMI prepared as described above. The plates were incubated at 35°C for 48 h. Growth inhibition was monitored by spectophotometric determination of optical density at 530 nm in a Molecular Devices SpectraMax^®^ PARADIGM^®^ microplate reader. The MIC was defined as the lowest concentration inhibiting 90% of cryptococcal growth. Sterility control wells were included in all plates. Similar protocols were used for determination of MICs for amphotericin B, fluconazole and benzimidazole derivatives. To evaluate whether fenbendazole was fungicidal or fungistatic, the cells were incubated at the MIC under the conditions described above, washed and plated on Sabouraud agar plates and incubated for 48 h at 30°C for determination of colony forming units (CFU). As preconized by the EUCAST, *Candida parapsilosis* and *C. krusei* strains (ATCC 22019 and ATCC 6258, respectively) were used as controls of the antifungal activity of amphotericin B and fluconazole.

### Checkerboard assay

The effects of the association of fenbendazole with amphotericin B on antifungal activity were evaluated by the checkerboard assay (40). The drug concentration ranges used in this assay corresponded to 0.007 - 0.5 ug/ml for amphotericin B and 0.003 - 1.536 μg/ml for fenbendazole. Drug solutions were prepared as previously described (12). Briefly, 50 μl of each of the amphotericin B concentrations was mixed with 50 μl of each of the fenbendazole solutions in microtiter plates. Vehicle concentration corresponded to 1% in all wells. Inoculation of fungal suspensions and evaluation of fungal growth followed the EUCAST protocol described in this section. The fractional inhibitory concentration index (FICI) was calculated according to the equation: ΣFICI = FICI (fenbendazole) + FICI (AmB), where the FICI was the ratio of the MIC of the combination with the MIC alone (41, 42).

### Cytotoxicity of fenbendazole

The cytotoxicity of fenbendazole to macrophages was determined using the Cytotox 96^®^ non-radioactive cytotoxicity assay kit (Promega, catalog number G1780), following the manufacturer’s recommendations. As recently described by our group (43), macrophages were used for cell viability assays due to their fundamental roles in the control and/or dissemination of cryptococci (27). RAW 264.7 macrophages (10^5^ cells/well in DMEM medium supplemented with 10% FBS were treated with 0.006 to 5.98 μg/ml fenbenzaole, alone or in combination with 0.06 μg/ml amphotericin B for 24 h at 37 °C in DMEM supplemented with 10% FBS. Cytotoxicity was inferred from the determination of the levels of lactate dehydrogenase activity in the medium. Control systems included vehicle-treated cells (viability control) and macrophages lysed with the lysis solution provided by the manufacturer (death control).

### Effects of fenbendazole on microtubule organization

Tubulin organization in *C. neoformans* and *C. gattii* was evaluated as described by Wang *et al*. (44) with minor modifications. Briefly, fungal cells were cultivated overnight in liquid Sabouraud medium at 30°C under shaking (200 rpm). These cultures had their optical density at 600 nm (OD_600_) adjusted to 0.2 in fresh, liquid Sabouraud, and then incubated at 30°C under shaking (200 rpm) until an OD_600_ value of approximately 0.8 was reached. Samples of 1 ml of these cell suspensions were washed with PBS and resuspended in fresh Sabouraud supplemented with 0.12 μg/ml fenbendazole (10 times concentrated MIC value) or DMSO (0.012 %). The suspensions were incubated for additional 90 min at 30°C with shaking (200 rpm) and washed with PBS. The cells were finally suspended in 1 ml of fresh Sabouraud supplemented with 0.5 μl of the Tubulin Tracker™ Green reagent (Thermo Fisher Scientific, catalog number T34075). The cells were incubated for 1 h at 37°C, washed with PBS and fixed with 2% paraformaldehyde in PBS. The samples were observed under a DMi8 fluorescence microscope (Leica). Images were recorded with the LasAF software.

### Effect of fenbendazole on capsular morphology

*C. neoformans* and *C. gattii* were grown overnight in liquid the Yeast Extract-Peptone-Dextrose (YPD) medium at 30°C and washed with PBS. Yeast suspensions had their density adjusted to 5 × 10^4^ cells/ml in a capsule induction medium (10% Sabouraud in 50 mM MOPS, pH 7.3) (45) supplemented with variable concentrations of fenbendazole. Control systems contained the vehicle alone. Capsule enlargement was induced at 37°C in a 5% CO_2_ atmosphere for 24 h. The cells were then washed with PBS and prepared for different microscopic analyses. For India ink counterstaining, paraformaldehyde (4%)-fixed cells were washed in PBS and mixed with India ink (1:1,v/v) for observation in a light microscope. Capsule dimensions were determined in digitalized imagens using the ImageJ software (46). For immunofluorescence, the paraformaldehyde-fixed cells were initially blocked with 1% bovine serum albumin (BSA) in PBS for 1 h at 37°C, following staining of cell wall chitin with 25 μM calcofluor white (Sigma, catalog number 18909) in PBS for 30 min at 37°C. The cells were washed 3 times with PBS and incubated for 1 h at 37°C with a monoclonal antibody (mAb 18B7, donated by Dr. Arturo Casadevall) raised to capsular glucuronoxylomannan (GXM) at 10 μg/ml in a PBS solution containing 1% BSA. The cells were washed with PBS and incubated with an Alexa 546-antibody conjugate (Invitrogen, catalog number A-11030) recognizing mouse immunoglobulin G (1 h, 25°C in the dark). Washed cells were analyzed in a Leica SP5 AOBS laser confocal microscope. For scanning electron microscopy (SEM), the cells were washed with PBS and fixed with 2.5% glutaraldehyde in 0.1 M sodium cacodylate buffer (pH 7.2) for 1 h at 25°C. The cells were washed with a 0.1 M sodium cacodylate buffer (pH 7.2) containing 0.2 M sucrose and 2 mM MgCl2. The cells were placed over 0.01% poly-L-lysine-coated coverslips and incubated for 30 min at 25°C. Adhered cells were then gradually dehydrated in ethanol (30, 50, and 70% for 5 min, and then 90% for 10 min, and 100% twice for 10 min). Immediately after dehydration, the cells were critical point dried (Leica EM CPD300), mounted on metallic bases and coated with a gold layer (Leica EM ACE200). The cells were visualized in a scanning electron microscope (JEOL JSM-6010 Plus/LA) operating at 5 keV.

### Intracellular antifungal activity

RAW 264.7 macrophages were suspended (5×10^5^ cells/ml) in DMEM supplemented with 10% FBS, and 200 μl of this suspension was transferred to each well of 96-wells plates for overnight incubation at 37°C in a 5% CO_2_ atmosphere. In the next day, *C. neoformans* or *C. gattii* (5 x 10^5^ cells/ml) were opsonized by incubation for 1 h at 37°C under a 5% CO_2_ atmosphere in DMEM containing 10% FBS and 5 μg/ml mAb 18B7. Macrophage cultures had their medium replaced with 200 μl of the above-described cell suspensions containing opsonized fungi and incubated for 2 h (37°C, 5% CO_2_). The systems were washed 3 times with PBS to remove unattached fungal cells and covered with 200 μL of DMEM containing 10% FBS, in addition to fenbendazole and/or amphotericin B (0.006 or 0.012 μg/ml fenbendazole; 0.06 or 0.5 μg/ml amphotericin B; 0.006 μg/ml fenbendazole combined with 0.06 μg/ml amphotericin B). Drug vehicle alone was used as a control. For determination of fungal killing, drug-containing systems were incubated for 24 h (37°C, 5% CO_2_) and washed 3 times with PBS. The macrophages were then lysed with 200 μl of cold, sterile water and the resulting suspensions were plated on solid Sabouraud medium. The systems were incubated at 30°C for 48 h and CFU were counted manually. For determination of intracellular proliferation and phagocytic escape of cryptococci, the plates containing infected macrophages were incubated at 37°C with 5% CO_2_ for 18 h in an Operetta High-Content Imaging System (PerkinElmer). During this 18 h incubation, images of each well were captured every 5 minutes with a 40x objective. Using the Harmony High Content Imaging and Analysis Software (PerkinElmer), movies of each well were prepared for analysis of intracellular proliferation rates (IPR) and vomocytic escape (38). Fifty cells were analyzed in each experimental condition. The IPR were calculated as the *t_18_/t_0_* ratio, where *t_18_* corresponded to the number of fungal cells inside the macrophages after 18 h and *t_0_* corresponded to the number of phagocyte-associated cryptococci at the beginning of the incubation. Vomocytosis was calculated as previously described (38) using 50 cells per experimental system. Three independent experiments were performed.

### Animal experimentation

For intranasal administration of fenbendazole, C57BL/6 mice (8 to 12-week-old) were used. The animals were kept with food and water *ad libitum* at the animal facility of the University of Brasília. The animals were placed in an isoflurane inhalation system (Bonther) for anesthesia and then challenged intratracheally with 25 μl of PBS containing 1×10^4^ yeasts of *C. neoformans* (H99 strain). After three days of infection, the animals were divided into groups with 5 individuals each. One group was treated intranasally with 20 μl of 1.25 μM fenbendazole. The other two groups were treated with PBS or amphotericin B (2 mg/kg) intraperitoneally as indicated in Figure 3. Mice were fed ad libitum and monitored every day for discomfort and signs of disease. Mice showing weight loss, lethargy, tremor, or inability to reach food or water were euthanized, and survival was counted until that day. At day 30 any surviving mice were euthanized. Euthanasia was performed with CO_2_ asphyxiation with 100% FiCO_2_ for 2 min, followed by cervical dislocation. The experiment was repeated under the same conditions and identical results were found. All treatments and experimental procedures were performed after approval by the Ethics Committee on Animal Use (CEUA) of the University of Brasília (UnBDoc no. 66729/2016) and according to the guidelines presented by the National Council for the Control of Experimentation Animal (CONCEA).

## Acknowledgements

We are thankful to Drs. Arturo Casadevall for donation of the 18B7 antibody and Charley Staats and Ane Garcia for the use of the *nopl6Δ* and *aim25Δ* mutants. We also appreciate the notation of the fluorescent strains by Dr. Robin May. M.L.R. is currently on leave from the position of Associate Professor at the Microbiology Institute of the Federal University of Rio de Janeiro, Brazil.

